# Neural flexibility in metabolic demand dynamics reveals sex-specific differences and supports cognition in late childhood

**DOI:** 10.1101/2025.09.24.678309

**Authors:** Aline Kotoski, Sir-Lord Wiafe, Vince D. Calhoun

**Affiliations:** Tri-Institutional Center for Translational Research in Neuroimaging and Data Science (TReNDS), Georgia State University, Atlanta, USA

**Keywords:** neural flexibility, sex differences, late childhood cognition, dynamic time warping, metabolic energy coordination

## Abstract

Dynamic coordination of metabolic demand across brain networks supports emerging cognitive abilities and may drive overall cognitive development, yet how these dynamics vary by sex and relate to cognition in late childhood remains unclear. Using resting-state fMRI from 2,000 healthy 9-to 11-year-olds in the ABCD study, we applied time-resolved dynamic time warping to quantify amplitude mismatches, a proxy of relative energy demand across brain intrinsic networks. Clustering revealed three recurring states: convergent (globally balanced), divergent (imbalanced), and mixed (intermediate). Females spent engaged more with the flexible mixed state, whereas males lingered longer in convergent and divergent states. Across the cohort, better performance on cognitive flexibility, processing speed, and long-term memory tasks correlated with greater overall time in the mixed state and with higher transition rates, but with shorter dwell in any single state. These findings indicate that neural flexibility, rather than prolonged stability, supports cognition during late childhood and that sex differences in dynamic energy coordination emerge well before adolescence.

## I. Introduction

Late childhood marks a critical stage of brain development characterized by dynamic changes in both structure and function [1], which support the development of complex cognitive processes. During this period, neural systems become increasingly specialized and interconnected, supporting improvements in attention, memory, and executive function [2]. These changes are supported by increased metabolic demand, as evidenced by rising cerebral blood flow and oxygen consumption throughout development [3]. Importantly, developmental trajectories may differ by sex, with males and females showing distinct patterns of brain maturation that are often reflected in differences in cognitive performance [4]. Understanding how sex and cognitive differences independently relate to dynamic patterns of energy demand provides important insight into the neural processes that shape brain function during late childhood. A key aspect of this maturation is the refinement of neural flexibility, the brain’s ability to fluidly transition between different network configurations to support ongoing cognitive demands [5]. Increasing evidence suggests that efficient reconfiguration of brain networks, rather than prolonged engagement in stable states, is a hallmark of healthy cognitive function [6, 7]. This dynamic interplay underscores the importance of analytical approaches that can capture transient, moment-to-moment fluctuations in neural processing.

Resting-state functional magnetic resonance imaging (rs-fMRI) offers a window into the brain’s intrinsic activity by capturing spontaneous fluctuations in the blood-oxygenation-level-dependent (BOLD) signal, which indirectly reflects ongoing neural processing and energy consumption [8]. However, traditional functional connectivity measures, such as Pearson correlation and sliding-window approaches, primarily quantify synchrony between brain regions and often overlook the amplitude dynamics embedded in the BOLD signal [8]. Because these methods typically normalize signal amplitudes [9], they are less effective at capturing transient mismatches in regional energy demands [10]. This limitation is particularly relevant during development, when neurovascular coupling and hemodynamic responses exhibit variability both across individuals and across neural circuits [11].

Recently, there has been growing interest in using dynamic time warping (DTW) as an alternative metric for functional connectivity in fMRI data [11-15]. Unlike conventional correlation-based approaches, DTW enables the detection of signal amplitude differences while warping temporal axis of the signals [16]. Building on this, a recent study developed a time-resolved DTW framework and discussed how it might map to metabolic energy demands from the fMRI data [8]. This method allows for a more nuanced investigation of how neural systems dynamically converge or diverge in their metabolic energy profiles, which may be particularly relevant during periods of intense brain maturation such as late childhood.

In this study, we apply time-resolved DTW to rs-fMRI data from children aged 9 to 11 years old, using the Adolescent Brain Cognitive Development (ABCD) study dataset. Our goal is to investigate how amplitude mismatches between resting-state brain networks, interpreted as metabolic energy demand differences, relate to individual differences in cognitive performance. By combining DTW-based metrics with standardized cognitive assessments, we aim to identify developmental signatures of neural efficiency and energy coordination that underlie cognitive function in childhood. Additionally, we examined sex differences in these dynamic brain patterns to understand whether males and females exhibit distinct trajectories of metabolic energy coordination.

## II. Methods

### A. DTW cumulative-window connectivity

Given two BOLD time series *x*(*t*) and *y*(*t*) of length *N*, we compute the DTW distance using a Sakoe–Chiba window [16] of half-width *w*. First, we define the local cost as:

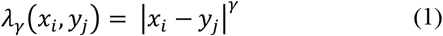

We use *γ* = 2 as we show in our previous study that higher *γ* ≥ 1.5 yields the most reliable parameter using a test-retest reliability test [8]. The cumulative cost matrix of the DTW algorithm *CM*_(*i,j*)_ for |*i* − *j*| ≤ *w* is built recursively as:

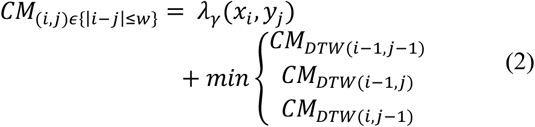

Once *CM* is complete, we extract the (*φ*_2_(*τ*), *φ*_3_(*τ*)) of length |*τ*| < *N* computed by backtracking from the *CM* matrix. From this path, define the time-resolved DTW distance:

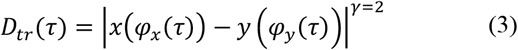

We then apply a piecewise cubic Hermite interpolating polynomial (PCHIP) to *D_tr_*(*τ*) so that it is resampled at integer time indices *i* = 1, …, *N*, yielding 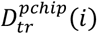.

Since a low-pass filter at 0.01 *Hz* is applied to BOLD signals, meaningful fluctuations occur over ∼100 *s*; more precisely, the −3*dB* cutoff corresponds to ∼88*s* [9, 17, 18]. We therefore choose a window size of ∼88*s* for the DTW algorithm.

### B. Recurring states estimation

Whole-brain time-resolved DTW across brain region pairs were clustered with standard *k*-means because of its widespread use in time-resolved fMRI analysis [19]. We implemented the *k*-means algorithm with cluster numbers ranging from 1 to 10, setting the maximum number of iterations to 10,000 and using 20 random initializations to ensure robust convergence. The city-block (Manhattan) distance metric was selected for its demonstrated robustness in handling high-dimensional data [20]. The optimal cluster number was determined as 3 using the elbow criterion.

### C. State dynamics

To capture individual temporal patterns from the k-means clustering results, we extracted three metrics. The mean dwell time (MDT) represents the average length of time a participant remains continuously in a given state, providing an index of state stability [21]. The fractional rate (FR) indicates the proportion of the total scan duration spent in each state, offering a measure of overall state prevalence [21]. Lastly, state-to-state transition probabilities capture the likelihood of switching from one state to another, shedding light on the directional dynamics of state changes over time [21].

### D. Group analysis & cognitive scores association

We assessed group differences between males and females, and associations with cognitive scores using generalized linear models that included age and scanning site as covariates. For cognitive scores, sex was included as a covariate. Multiple comparisons were controlled with the Benjamini–Hochberg false discovery rate (FDR) procedure.

### E. Data

This study used rs-fMRI and cognitive data from the ABCD study, a large-scale, multisite longitudinal project designed to track brain and behavioral development from childhood through adolescence [22, 23]. A total of 2,000 children (1,011 males) aged between 8 years and 11 months and 11 years were included in this analysis. The mean age of the sample was 9 years and 10 months (SD = 7 months). On average, males were 9 years and 11 months old (SD = 7 months), and females were 9 years and 10 months old (SD = 7 months).Only one scan per subject was used, with no repeated measures or longitudinal data analyzed in this study.

Head motion in the ABCD resting-state fMRI data is carefully addressed through multiple preprocessing steps. To mitigate motion artifacts, volumes exceeding a framewise displacement (FD) threshold of 0.3 mm, along with one preceding and two following frames, are censored during motion scrubbing [22]. Participants with excessive motion are excluded from further analysis [22]. In addition, nuisance regression incorporates 24 motion parameters (translations, rotations, their derivatives, and squared terms), along with physiological regressors derived from white matter and cerebrospinal fluid signals, to reduce residual motion and physiological noise [22]. These procedures collectively aim to ensure that the resting-state fMRI signals predominantly reflect intrinsic neural activity rather than motion-related or physiological confounds [22].

### F. fMRI parameters & processing

Data were collected on three types of 3T scanners (Siemens Prisma, General Electric 750 and Philips) all with a standard adult-size head coil. The resting-state EPI sequence used the following parameters: TR = 800 ms, TE = 30 ms, 60 slices, flip angle = 520, matrix size = 90 x 90, FOV = 216 x 216 mm, resolution = 2.4 x 2.4 x 2.4 mm. The full details of the imaging acquisition protocol are described in [22].

All volumes underwent conventional preprocessing steps, including slice-timing correction, rigid-body motion correction, spatial normalization to MNI space, and spatial smoothing with a 6 mm full-width at half-maximum (FWHM) Gaussian kernel [24]. Spatially independent components were then identified using the NeuroMark 1.0 ICA pipeline [25], resulting in 53 intrinsic connectivity networks (ICNs) consistently observed across all participants. The corresponding ICN time courses were subsequently processed to remove linear trends, suppress outlier spikes, and bandpass filtered using a 7th-order Butterworth filter with a frequency range of 0.01–0.15 Hz. This filter configuration, optimized via MATLAB’s *Buttord* function, ensured a passband ripple below 3 dB and at least 30 dB attenuation in the stopband. Finally, the filtered time series were normalized using z-score transformation.

### G. Cognitive scores

To evaluate individual differences in cognitive performance, we used a broad set of age-corrected scores from the NIH Toolbox Cognition Battery [26] and the Rey Auditory Verbal Learning Test (RAVLT) [27], both administered as part of the ABCD baseline assessment. These measures were selected to capture a range of cognitive domains relevant to childhood brain development, including executive function, memory, language, and general intellectual ability.

From the NIH Toolbox, we included the following age-corrected scores:

- Flanker Inhibitory Control and Attention Test.
- List Sorting Working Memory Test.
- Dimensional Change Card Sort (DCCS) Test.
- Pattern Comparison Processing Speed Test.
- Picture Vocabulary Test.
- Oral Reading Recognition Test.
- Fluid Cognition Composite Score, Crystallized Cognition Composite Score, and Total Cognition Composite Score.

In addition to these age-adjusted scores, we also included the theta scores for the reading skill, and language ability. For the RAVLT, we included the short-delay, long-delay tasks, and total correct responses. More information about these cognitive assessments, including task descriptions, scoring procedures, and validation details, is available in the NIH Toolbox and ABCD study documentation [22, 26, 28].

## III. Results

### A. Group differences between males and females

We derived 3 recurring states of the time-resolved DTW metric suggesting 3 global states of metabolic energy demands. To ensure positive values represent convergence and negative values indicate divergence, we subtracted the median of all clusters and multiplied the distribution by −1 (Fig. 1a). We classified the clusters as: state 1 – convergent state, state 3 – divergent state, and state 2 – a mixed state that shows neither strong convergence nor divergence. These state-specific matrices are visualized in Figure 1b.

**Figure 1.**
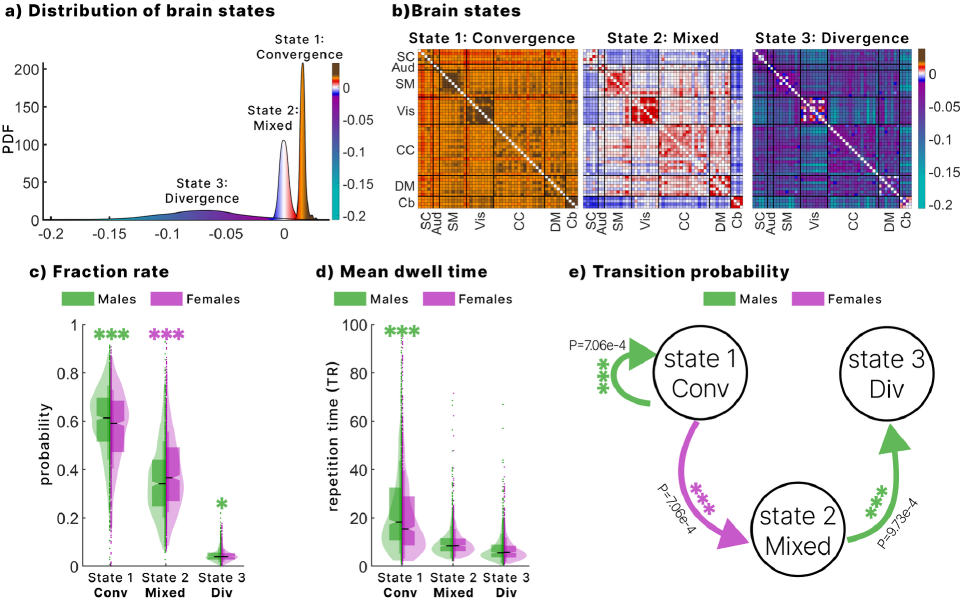
Group differences between males and females: Recurring brain states were identified by applying k-means clustering to time-resolved DTW metric. To interpret the directionality of network dynamics, the median across all three cluster centroids was subtracted and the result multiplied by –1, such that positive values indicate homogeneous amplitude disparity (convergence) and negative values indicate heterogenous amplitude disparity (divergence). a. Distribution of the recurring brain states b. The recurring brain state matrices c. Fraction rate in each state across sex groups, highlighting increased fraction rate in convergence and divergence state for males and increased mixed state for females. d. Mean dwell time in each state across sex groups, highlighting increased mean dwell time in convergence state for males. e. Transition matrix showing differences in probability of males and female transition between states. One asterisk (*) indicates FDR-corrected p-values between 0.05 and 0.01, two asterisks (**) indicate p-values between 0.01 and 0.001, and three asterisks (***) indicate p-values below 0.001.

We examined sex differences in several metrics derived from the time-resolved DTW analysis, including mean dwell time, fractional rate, and state transition probabilities. Females showed greater engagement with the mixed state, as indicated by a higher FR, while males exhibited greater engagement with both the convergent and divergent states (Fig. 1c). Additionally, males demonstrated significantly longer mean dwell times in the convergent state, suggesting a stronger tendency to remain in that state compared to females (Fig. 1d). Regarding state transition probabilities, females were more likely to transition from the convergent state to the mixed state (Fig. 1e). In contrast, males were more likely to exhibit self-transitions within the convergent state and to transition from the mixed state to the divergent state, relative to females.

### B. Cognitive scores associations

To assess how predictive the time-resolved DTW metric were of the cognitive scores across all participants, performed generalized linear models across all the three state dynamic metrics. Several cognitive scores showed significant associations; however, we focus our results on three scores that were significantly associated across all the three DTW-derived metrics: MDT, FR, and state transition probabilities. These three scores include:

- The Dimensional Change Card Sort (DCCS) test: evaluates cognitive flexibility by requiring participants to sort cards according to shifting rules.
- The Pattern Comparison test: measures visual processing speed by asking participants to quickly judge whether two visual patterns are the same.
- The RAVLT long-delay recall test: evaluated long-term verbal memory by measuring delayed recall of the original word list.

State transition probabilities (Fig. 2a) revealed meaningful associations with cognitive performance. A higher probability of remaining in the convergent state was significantly associated with lower scores across all three cognitive measures. Conversely, increased transitions from the mixed state to the convergent state were linked to better performance on the DCCS and RAVLT long-delay tests, while transitions from the mixed to the divergent state were associated with poorer performance on these same measures. Specifically, for the DCCS test, better scores were also associated with more frequent transitions from the divergent to the mixed state, whereas worse performance correlated with a higher probability of remaining in the divergent state. For the RAVLT long-delay test, stronger transitions from the convergent to the mixed state were also linked to higher scores. Together, these findings suggest that sustained occupancy in any single state, particularly the convergent or divergent states, is associated with poorer cognitive performance, while dynamic transitions, especially those avoiding the divergent state, are generally predictive of better outcomes.

**Figure 2.**
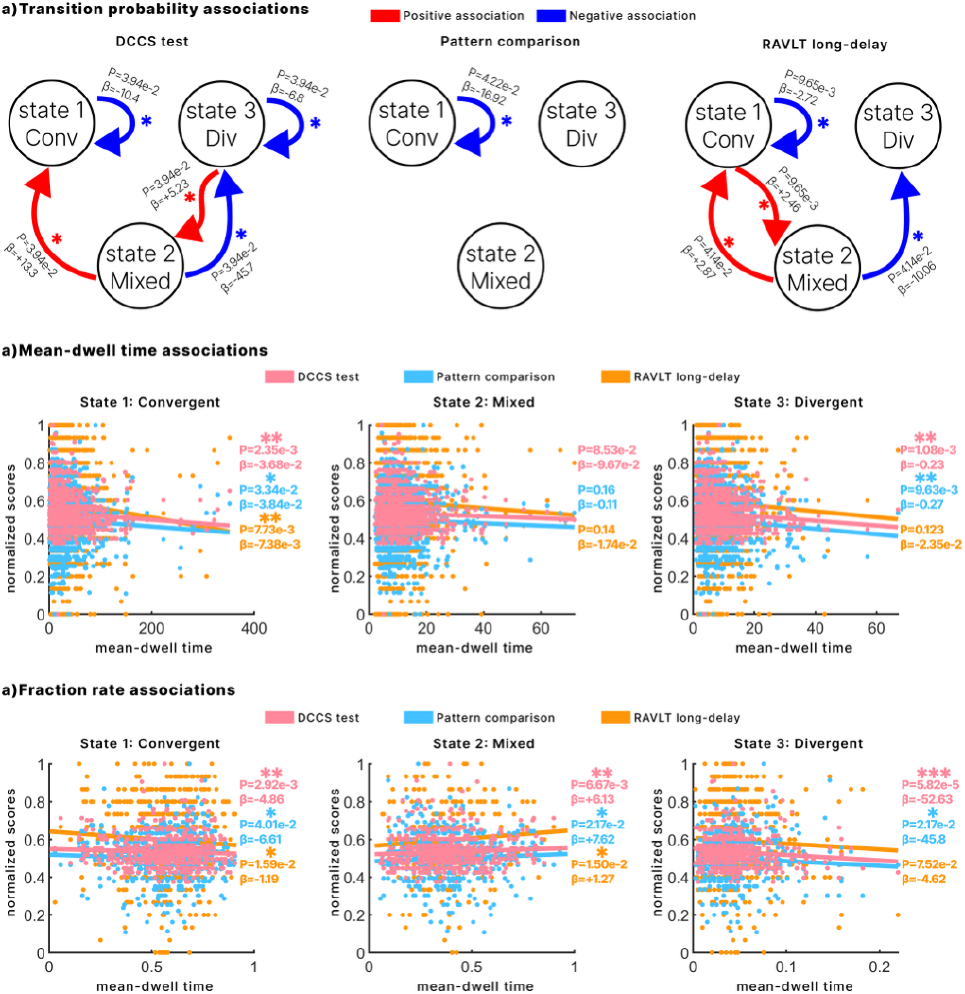
illustrates the association of 3 cognitive tests with the three derived state dynamic metrics from the time-resolved DTW metric using a generalized linear model including age, sex, and site as covariates (FDR-corrected). a. illustrates the state transitions and their association with the 3 cognitive tests with a blue color indicating a negative association and a red indicating a positive association. b. Associations between mean-dwell time and the three cognitive scores across all three states. c. Associations between fraction rate and the three cognitive scores across all three states. All the three scores are normalized between 0 and 1 for visualization purposes. One asterisk (*) indicates FDR-corrected p-values between 0.05 and 0.01, two asterisks (**) indicate p-values between 0.01 and 0.001, and three asterisks (***) indicate p-values below 0.001.

For the fractional rate, greater total engagement in the mixed state was significantly associated with higher performance across all three cognitive measures. In contrast, greater engagement in the convergent and divergent states was consistently linked to lower cognitive scores. Regarding mean dwell time, longer persistence in the convergent and divergent states was significantly associated with poorer cognitive outcomes. Although dwell time in the mixed state did not show statistically significant associations with cognitive performance (p = 0.085, 0.15, and 0.14, respectively), the trend still indicated that greater persistence in this state was related to lower scores.

## IV. Discussion

Our findings reveal distinct sex differences in dynamic patterns of power imbalance across resting-state brain networks during late childhood. Using time-resolved DTW and k-means clustering, we identified three recurring states; labeled as convergent, divergent and mixed. The convergent state reflects a global brain condition in which blood flow is relatively uniform across regions, suggesting a homogeneous distribution of metabolic demand [8]. In contrast, the divergent state indicates pronounced heterogeneity in regional blood flow [8]. The mixed state represents an intermediate condition with no strong bias toward convergence or divergence.

Both male and female participants in our healthy ABCD cohort spent the greatest proportion of time in the convergent state, a globally balanced, low-cost configuration as illustrated by the fraction rate in Fig. 1c. This predominance is fully consistent with earlier dynamic connectivity work showing that healthy brains dwell longest in energy-balanced states whereas clinical populations (e.g., schizophrenia) occupy them far less frequently [8, 19].

Through transition probabilities and fraction rate, females exhibited significantly higher engagement with the mixed state, a metastable “critical” pattern that hovers between order and disorder and can re-configure at very low energetic cost. Two lines of prior evidence make this plausible. First, key maturational milestones—fronto-parietal myelination, synaptic pruning, and neurovascular fine-tuning—reach their plateaus roughly a year earlier in girls [29, 30], which should reduce the metabolic burden of switching network configurations and render the mixed state an efficient default. Second, dynamic-connectivity studies show that adolescent and adult females enter and leave resting-state configurations more frequently than males; the mixed state, by definition an ultra-short alternation between convergent- and divergent-like layouts, neatly captures this higher transition propensity [31]. Altogether, higher female engagement of the mixed state likely reflects a combination of earlier structural maturation and faster functional switching.

Through analyses of fractional rate, mean dwell time, and transition probabilities, we found that males preferentially engaged the convergent state and, via both fractional rate and transition matrices, engaged the divergent state more often compared to females. Their significantly longer dwell times in convergent indicate that once male brains settle into this globally synchronized, energy-balanced configuration they remain there for extended periods, implying slower disengagement and reduced flexibility in reallocating metabolic resources. This pattern mirrors dynamic connectivity work in the Human Connectome Project, where males persisted longer within salience-network states whereas females cycled more rapidly among configurations [31]. It also dovetails with longitudinal neurodevelopmental evidence showing that structural and neurovascular maturation— cortical thinning, fronto-parietal myelination, and strengthening of cerebral blood-flow/BOLD coupling—peaks later in boys than in girls [32]. Our results suggest that late-childhood males rely on a more rigid or bimodal energetic strategy: prolonged sojourns in a stable, low-cost convergent mode punctuated by comparatively frequent but metabolically costly excursions into divergent dynamics. Our finding that males and females exhibit distinct dynamic functional styles is convergent with our prior work. Using the same ABCD cohort, we previously showed that the sexes also differ in their patterns of dynamic coupling between brain structure (sMRI) and functional network connectivity (dFNC), with each sex demonstrating stronger coupling in a unique set of brain networks [33].

Across participants, better performance on the three behavioral measures—cognitive flexibility (DCCS), visual processing speed (Pattern Comparison) and long-term memory (RAVLT long-delay)—tracked a *higher fractional rate* in the mixed state and a *lower fractional rate* in both convergent and divergent states. The mixed state sits on the ridgeline between order and disorder, so revisiting it often gives the system many low-cost launch points for re-organizing its network layout. Large ABCD samples likewise show that children who excel on composite cognition spend proportionally more time in weakly-connected, metastable configurations and less time in globally coherent (or strongly segregated) ones [34]. The principle that cognitive proficiency is supported by a specific pattern of dynamic brain organization is further supported by our multimodal work in the same cohort. Using a different approach that links brain structure and function, we previously demonstrated that higher reading ability was specifically associated with stronger structure-function coupling in higher-order cognitive networks, such as fronto-parietal circuits, and weaker coupling in primary visual areas [23]. Similar benefits of network flexibility have been reported in adults for general intelligence and creativity [35].

Mean-dwell time told the complementary story: *lingering* in any one state—Convergent, Divergent or Mixed—was associated with poorer scores (the Mixed effect fell just short of significance but followed the same direction). Previous developmental work shows that children with longer dwell times in highly coherent or globally disconnected states display weaker executive skills [36], and theoretical accounts argue that cognitive growth depends on the ability to exit an attractor quickly rather than on the attractor’s absolute prevalence [37].

Putting the two metrics together, our results suggest that spending a lot of total time in mixed is advantageous only when that time is achieved through many brief visits, not through “sticky” persistence. In other words, the mixed state appears to act as a flexible hub that propels transitions [8]; frequent returns to it boost cognition, but getting trapped there, like getting trapped anywhere, dampens performance.

Transition probability analyses sharpen this picture of “flexibility beats stability.” Children who moved often from convergent to mixed, mixed to convergent, or divergent to mixed were associated with the higher scores on the cognitive tests, implying that the mixed state acts as a low-energy relay that lets the system reshuffle its network layout on demand [8, 38]. The only negative transition effect appeared for mixed to divergent on the RAVLT, reinforcing the view that engagement with the metabolically costly divergent state hamper cognition [8, 39]. Likewise, a high self-transition probability in convergent, essentially staying put in that “attractor”, was uniformly linked to poorer performance across all three measures, mirroring findings that long dwell in globally synchronous states predicts weaker executive skills in children [34]. Together with prior work showing that greater transition entropy and neural dynamics closer to criticality predict higher fluid intelligence and working-memory capacity in adolescents and adults [38, 40], these results suggest that, in late childhood, cognitive efficiency depends less on occupying any particular state and more on the brain’s ability to keep cycling among states without getting “stuck.”

Taken together, our findings provide compelling evidence for the central role of neural flexibility in supporting cognitive maturation during late childhood. This flexibility manifests in two complementary ways. First, at the level of temporal dynamics, cognitive proficiency is associated not with stability but with the brain’s agility, its capacity for rapid state transitions and brief dwell times. Second, at the level of state architecture, the mixed state appears to function as a critical hub. Our results suggest that for the developing brain, frequent access to this flexible hub state is more advantageous for cognition than prolonged occupancy in more rigid configurations.

Despite the strengths of our approach, a few limitations should be acknowledged. The cross-sectional design of this study limits developmental inferences, longitudinal analyses will be essential to determine whether the identified dynamic state patterns predict later cognitive outcomes. While we interpret our findings in the context of neural efficiency and energy, BOLD fMRI is an indirect metabolic proxy. Future work integrating complementary modalities such as arterial spin labeling or FDG-PET will be important to validate and refine energy-related interpretations. Although we selected the number of clusters using the elbow criterion, consistent with the original study that introduced the time-resolved DTW method, we did not systematically explore how varying the number of clusters or using alternative distance metrics might affect state definitions or their cognitive associations. Assessing the robustness of findings across different clustering solutions is an important direction for future work. While we covaried for scanner manufacturer to account for site effects, we did not perform a detailed exploration of how specific hardware, such as differences in head coil configurations across sites, might systematically bias our amplitude-based metrics. Future technical studies will be needed to assess the robustness of these dynamic states to such equipment-related signal variations. Additionally, while test– retest reliability of the time-resolved DTW approach was established in prior studies, we did not replicate this analysis in our dataset, which may limit generalizability. Finally, although we observed associations across multiple cognitive domains, we focused on the most consistently associated scores to highlight core findings. Future analyses will expand on these results by examining the full cognitive battery to better characterize domain-specific and domain-general contributions. It is also important to acknowledge that while our results are statistically significant, the observed effect sizes are modest. This is consistent with the broader literature on brain-behavior associations in large cohorts, and it suggests that while these dynamic states provide meaningful insight into the neural basis of cognition, their value for making practical, individual-level predictions will depend on their integration into more comprehensive, multivariate models in future work.

In summary, late-childhood brains appear to profit from metabolic agility. Females engage more time in the mixed or “critical” state and switch among states more often, signaling a flexible energy-coordination style. Males gravitate toward longer, more rigid sojourns in the energy-balanced convergent state and, intermittently, the costly divergent state. Crucially, better performance on cognitive flexibility, processing speed and long-term memory tasks align with greater overall time in the mixed state and with brisk shuttling among states, whereas prolonged residence in any single state predicts poorer scores. These patterns indicate that, during late childhood, it is the capacity to *move*, rather than to *stay*, that underpins efficient cognition, highlighting dynamic metabolic flexibility as a key developmental asset.

## V. Conclusion

This study demonstrates that time-resolved dynamic time warping applied to rs-fMRI can reveal meaningful patterns of energy coordination across brain networks during late childhood. Time-resolved DTW of resting-state fMRI revealed three recurring energy coordination states— convergent, divergent, and mixed—whose usage differs by sex and predicts cognition. Females favored the flexible mixed state and switched among states more often, whereas boys engaged more in convergent and divergent configurations. Across all children, better cognitive flexibility, processing speed, and memory aligned with more total time in the mixed state and with frequent state switching, while extended dwell in convergent or divergent predicted poorer scores. Thus, dynamic flexibility, rather than stability, appears to underpin efficient brain function during late childhood and follows sex-specific developmental trajectories.

## Acknowledgment

This work was supported in part by the NIH R01MH123610 and NSF 2112455.

## Notes

### Competing Interest Statement

The authors have declared no competing interest.

## References

[1] K. L. Mills, A. L. Goddings, L. S. Clasen, J. N. Giedd, and S. J. Blakemore, “The developmental mismatch in structural brain maturation during adolescence,” Dev Neurosci, vol. 36, no. 3-4, pp. 147–60, 2014, doi: 10.1159/000362328.

[2] G. L. Baum et al., “Development of structure–function coupling in human brain networks during youth,” Proceedings of the National Academy of Sciences, vol. 117, no. 1, pp. 771–778, 2020.

[3] T. Takahashi, R. Shirane, S. Sato, and T. Yoshimoto, “Developmental changes of cerebral blood flow and oxygen metabolism in children,” AJNR Am J Neuroradiol, vol. 20, no. 5, pp. 917–22, May 1999. [Online]. Available: https://www.ncbi.nlm.nih.gov/pubmed/10369366.

[4] H. C. Brenhouse and S. L. Andersen, “Developmental trajectories during adolescence in males and females: a cross-species understanding of underlying brain changes,” Neurosci Biobehav Rev, vol. 35, no. 8, pp. 1687–703, Aug 2011, doi: 10.1016/j.neubiorev.2011.04.013.

[5] D. S. Bassett, N. F. Wymbs, M. A. Porter, P. J. Mucha, J. M. Carlson, and S. T. Grafton, “Dynamic reconfiguration of human brain networks during learning,” Proceedings of the National Academy of Sciences, vol. 108, no. 18, pp. 7641–7646, 2011.

[6] J. M. Shine et al., “The Dynamics of Functional Brain Networks: Integrated Network States during Cognitive Task Performance,” Neuron, vol. 92, no. 2, pp. 544–554, Oct 19 2016, doi: 10.1016/j.neuron.2016.09.018.

[7] V. D. Calhoun, R. Miller, G. Pearlson, and T. Adalý, “The chronnectome: time-varying connectivity networks as the next frontier in fMRI data discovery,” Neuron, vol. 84, no. 2, pp. 262–274, 2014.

[8] S.-L. Wiafe et al., “Mapping Dynamic Metabolic Energy Distribution in Brain Networks using fMRI: A Novel Dynamic Time Warping Framework,” bioRxiv, p. 2025.03.20.644399, 2025.

[9] S. L. Wiafe, N. O. Asante, V. D. Calhoun, and A. Faghiri, “Studying time-resolved functional connectivity via communication theory: on the complementary nature of phase synchronization and sliding window Pearson correlation,” bioRxiv, Nov 22 2024, doi: 10.1101/2024.06.12.598720.

[10] S. L. Wiafe et al., “Mapping Dynamic Metabolic Energy Distribution in Brain Networks using fMRI: A Novel Dynamic Time Warping Framework,” bioRxiv, Mar 21 2025, doi: 10.1101/2025.03.20.644399.

[11] S.-L. Wiafe, A. Faghiri, Z. Fu, R. Miller, A. Preda, and V. D. Calhoun, “The dynamics of dynamic time warping in fMRI data: a method to capture inter-network stretching and shrinking via warp elasticity,” Imaging Neuroscience, vol. 2, pp. 1–23, 2024.

[12] S.-L. Wiafe, S. Kinsey, A. Iraji, R. Miller, and V. D. Calhoun, “Normalized Dynamic Time Warping Increases Sensitivity In Differentiating Functional Network Connectivity In Schizophrenia,” bioRxiv, p. 2024.10.31.621415, 2024.

[13] R. J. Meszlenyi, P. Hermann, K. Buza, V. Gal, and Z. Vidnyanszky, “Resting State fMRI Functional Connectivity Analysis Using Dynamic Time Warping,” Front Neurosci, vol. 11, p. 75, 2017, doi: 10.3389/fnins.2017.00075.

[14] L. Wiafe, A. Faghiri, Z. Fu, R. Miller, and V. D. Calhoun, “Capturing Stretching and Shrinking of Inter-Network Temporal Coupling in FMRI Via WARP Elasticity,” in 2024 IEEE International Symposium on Biomedical Imaging (ISBI), 2024: IEEE, pp. 1–4.

[15] A. C. Linke et al., “Dynamic time warping outperforms Pearson correlation in detecting atypical functional connectivity in autism spectrum disorders,” Neuroimage, vol. 223, p. 117383, Dec 2020, doi: 10.1016/j.neuroimage.2020.117383.

[16] K. K. Paliwal, A. Agarwal, and S. S. Sinha, “A modification over Sakoe and Chiba’s dynamic time warping algorithm for isolated word recognition,” Signal Processing, vol. 4, no. 4, pp. 329–333, 1982.

[17] A. Faghiri, A. Iraji, E. Damaraju, J. Turner, and V. D. Calhoun, “A unified approach for characterizing static/dynamic connectivity frequency profiles using filter banks,” Netw Neurosci, vol. 5, no. 1, pp. 56–82, 2021, doi: 10.1162/netn_a_00155.

[18] A. Faghiri et al., “Frequency modulation increases the specificity of time-resolved connectivity: A resting-state fMRI study,” Netw Neurosci, vol. 8, no. 3, pp. 734–761, 2024, doi: 10.1162/netn_a_00372.

[19] E. Damaraju et al., “Dynamic functional connectivity analysis reveals transient states of dysconnectivity in schizophrenia,” Neuroimage Clin, vol. 5, pp. 298–308, 2014, doi: 10.1016/j.nicl.2014.07.003.

[20] C. C. Aggarwal, A. Hinneburg, and D. A. Keim, “On the surprising behavior of distance metrics in high dimensional space,” in International conference on database theory, 2001: Springer, pp. 420–434.

[21] A. Iraji, A. Faghiri, N. Lewis, Z. Fu, S. Rachakonda, and V. D. Calhoun, “Tools of the trade: estimating time-varying connectivity patterns from fMRI data,” Soc Cogn Affect Neurosci, vol. 16, no. 8, pp. 849–874, Aug 5 2021, doi: 10.1093/scan/nsaa114.

[22] B. J. Casey et al., “The adolescent brain cognitive development (ABCD) study: imaging acquisition across 21 sites,” Developmental cognitive neuroscience, vol. 32, pp. 43–54, 2018.

[23] A. Kotoski, J. Liu, R. Morris, and V. Calhoun, “Inter-modality source coupling: a fully-automated whole-brain data-driven structure-function fingerprint shows replicable links to reading in a large-scale (N∼ 8K) analysis,” bioRxiv, p. 2024.03.07.583896, 2024.

[24] W. D. Penny, K. J. Friston, J. T. Ashburner, S. J. Kiebel, and T. E. Nichols, Statistical parametric mapping: the analysis of functional brain images. Elsevier, 2011.

[25] Y. Du et al., “NeuroMark: An automated and adaptive ICA based pipeline to identify reproducible fMRI markers of brain disorders,” Neuroimage Clin, vol. 28, p. 102375, 2020, doi: 10.1016/j.nicl.2020.102375.

[26] S. Weintraub et al., “Cognition assessment using the NIH Toolbox,” Neurology, vol. 80, no. 11_supplement_3, pp. S54–S64, 2013.

[27] A. Rey, “L’examen clinique en psychologie,” 1958.

[28] S. G. Heeringa and P. A. Berglund, “A guide for population-based analysis of the Adolescent Brain Cognitive Development (ABCD) Study baseline data,” BioRxiv, p. 2020.02.10.942011, 2020.

[29] R. K. Lenroot and J. N. Giedd, “Sex differences in the adolescent brain,” Brain Cogn, vol. 72, no. 1, pp. 46–55, Feb 2010, doi: 10.1016/j.bandc.2009.10.008.

[30] Z. Chen, H. Zhang, P. A. Yushkevich, M. Liu, and C. Beaulieu, “Maturation Along White Matter Tracts in Human Brain Using a Diffusion Tensor Surface Model Tract-Specific Analysis,” Front Neuroanat, vol. 10, p. 9, 2016, doi: 10.3389/fnana.2016.00009.

[31] L. Murray, J. M. Maurer, A. L. Peechatka, B. B. Frederick, R. H. Kaiser, and A. C. Janes, “Sex differences in functional network dynamics observed using coactivation pattern analysis,” Cogn Neurosci, vol. 12, no. 3-4, pp. 120–130, Jul-Oct 2021, doi: 10.1080/17588928.2021.1880383.

[32] J. E. Bramen et al., “Puberty influences medial temporal lobe and cortical gray matter maturation differently in boys than girls matched for sexual maturity,” Cerebral cortex, vol. 21, no. 3, pp. 636–646, 2011.

[33] A. Kotoski, S.-L. Wiafe, J. Stephen, Y.-P. Wang, T. Wilson, and V. Calhoun, “Dynamic Inter-Modality Source Coupling Reveals Sex Differences in Children based on Brain Structural-Functional Network Connectivity: A Multimodal MRI Study of the ABCD Dataset,” bioRxiv, p. 2025.07.23.666366, 2025.

[34] Z. Fu, J. Sui, A. Iraji, J. Liu, and V. Calhoun, “Cognitive and Psychiatric Relevance of Dynamic Functional Connectivity States in a Large (N>10,000) Children Population,” Res Sq, Jan 12 2024, doi: 10.21203/rs.3.rs-3586731/v1.

[35] T. A. Sassenberg, A. Safron, and C. G. DeYoung, “Stable individual differences from dynamic patterns of function: brain network flexibility predicts openness/intellect, intelligence, and psychoticism,” Cereb Cortex, vol. 34, no. 9, Sep 3 2024, doi: 10.1093/cercor/bhae391.

[36] H. A. Marusak et al., “Dynamic functional connectivity of neurocognitive networks in children,” Hum Brain Mapp, vol. 38, no. 1, pp. 97–108, Jan 2017, doi: 10.1002/hbm.23346.

[37] J. D. Medaglia et al., “Flexible traversal through diverse brain states underlies executive function in normative neurodevelopment,” arXiv preprint arXiv:1510.08780, 2015.

[38] X. Xin, J. Yu, and X. Gao, “The brain entropy dynamics in resting state,” Front Neurosci, vol. 18, p. 1352409, 2024, doi: 10.3389/fnins.2024.1352409.

[39] Y. Guo et al., “Dynamic functional connectivity changes associated with decreased memory performance in betel quid dependence,” Addict Biol, vol. 28, no. 10, p. e13329, Oct 2023, doi: 10.1111/adb.13329.

[40] L. Xu, J. Feng, and L. Yu, “Avalanche criticality in individuals, fluid intelligence, and working memory,” Hum Brain Mapp, vol. 43, no. 8, pp. 2534–2553, Jun 1 2022, doi: 10.1002/hbm.25802.

